# Hearing loss is associated with delayed neural responses to continuous speech

**DOI:** 10.1101/2021.01.21.427550

**Authors:** Marlies Gillis, Lien Decruy, Jonas Vanthornhout, Tom Francart

**Affiliations:** KU Leuven, Department of Neurosciences, ExpORL, 3000 Leuven, Belgium; Institute for Systems Research, University of Maryland, College Park, MD 20740, USA

**Keywords:** neural tracking, hearing loss, speech, EEG

## Abstract

We investigated the impact of hearing loss on the neural processing of speech. Using a forward modeling approach, we compared the neural responses to continuous speech of 14 adults with sensorineural hearing loss with those of age-matched normal-hearing peers.

Compared to their normal-hearing peers, hearing-impaired listeners had increased neural tracking and delayed neural responses to continuous speech in quiet. The latency also increased with the degree of hearing loss. As speech understanding decreased, neural tracking decreased in both populations; however, a significantly different trend was observed for the latency of the neural responses. For normal-hearing listeners, the latency increased with increasing background noise level. However, for hearing-impaired listeners, this increase was not observed.

Our results support the idea that the neural response latency indicates the efficiency of neural speech processing. Hearing-impaired listeners process speech in silence less efficiently than normal-hearing listeners. Our results suggest that this reduction in neural speech processing efficiency is a gradual effect which occurs as hearing deteriorates. Moreover, the efficiency of neural speech processing in hearing-impaired listeners is already at its lowest level when listening to speech in quiet, while normal-hearing listeners show a further decrease in efficiency when the noise level increases.

From our results, it is apparent that sound amplification does not solve hearing loss. Even when listing to speech in silence at a comfortable loudness, hearing-impaired listeners process speech less efficiently.

## Introduction

It is widely known that hearing loss alters the brain (Eggermont, 2017; Peelle and Wingfield, 2016). To study the functional neural changes, several studies focussed on cortical auditory evoked potentials (CAEP) using electroencephalography (EEG). CAEPs reflect the cortical responses evoked by repetitions of simple sounds such as syllables, tone pips, or clicks. These neural responses reflect sound detection and/or discrimination by the brain. The CAEP-response is characterized by a first positive peak (P1) around 50 ms, a first negative peak (N1) around 100 ms and a later positive peak (P2) around 180 ms (Burkard et al., 2007). Harkrider et al. (2009) and Campbell and Sharma (2013) reported increased P2-latencies in hearing impaired listeners(HI listeners) compared to normal hearing listeners (NH listeners). Interestingly, Campbell and Sharma (2013) reported that P2-latency was also correlated with the person’s speech perception ability in noise: a poorer perception of speech in noise is associated with a longer P2-latency. Although changes in latency are often not reported, in most studies HI listeners showed increased amplitudes compared to NH listeners (Tremblay et al., 2003; Harkrider et al., 2006; Bertoli et al., 2011; Alain, 2014; Maamor and Billings, 2017) while Billings et al. (2015) and Koerner and Zhang (2018) did not observe differences between these two populations. However, other studies attributed differences in neural response to the audibility of the stimulus: although presented at the same sound intensity, the sound might be less audible for HI listeners than for NH listeners or differently, when the sound is presented at a higher intensity for HI listeners, the neural differences between the two populations might be an effect of stimulus intensity rather than the impact of the hearing loss (Oates et al., 2002; Van Dun et al., 2016; McClannahan et al., 2019).

No consensus has been reached on the impact of hearing loss on the P1-N1-P2-complex. This may be due to the complexity of hearing-loss-related research: age and hearing loss are difficult to disentangle as HI listeners are often older. Moreover, when using simple sound stimuli, attention, motivation and task instructions might confound the study results and explain the variability between the findings of above-mentioned studies. In this study, participants listen to continuous speech. Continuous speech is closer to real-life situations than simple sounds or clicks. As understanding speech requires more in-depth neural processing of the stimulus, we believe that the use of continuous speech as a stimulus is essential to investigate the neural changes underlying speech understanding deficits in HI listeners.

When listening to continuous speech, neural responses time-lock to specific speech characteristics. This phenomenon is called neural tracking, i.e. the brain tracks specific characteristics of the speech (for a review, see, e.g., Brodbeck and Simon, 2020). Neural tracking can be derived from a forward or backward modeling approach. Most of the neural tracking studies focus on a measure of neural tracking derived from a backward modeling approach: a linear model fitted to reconstruct the acoustic representation, e.g., the envelope, from the measured EEG responses. This approach results in a reconstruction accuracy representing how well the acoustic representation is reconstructed from the EEG responses. With the forward modeling approach, a linear model is fitted to predict the EEG responses from the acoustic representation. Such a forward modeling approach yields a prediction accuracy, i.e., to what extent the predicted EEG responses correlate with the actual EEG responses, and a temporal response function (TRF). Such a TRF characterizes the way the brain responds to speech, as a function of measurement electrode and time delay, and therefore, gives unique temporal and spatial information.

Often neural tracking is studied with acoustic properties of the speech, like the envelope or spectrogram, and by using a backward modeling approach (Aiken and Picton, 2008; Ding and Simon, 2012b). Interestingly, in NH listeners, neural tracking increases when the speech is better understood (Ding and Simon, 2012a;Horton et al., 2014; O’Sullivan et al., 2015; Das et al., 2016; Etard and Reichenbach, 2019; Iotzov and Parra,2019; Vanthornhout et al., 2018; Lesenfants et al., 2019). A limited number of studies has been conducted to study the effect of hearing loss on neural tracking of continuous speech. These studies investigated the effect of hearing loss in a two-talker scenario: an attended speaker and an ignored one (Petersen et al., 2017;Mirkovic et al., 2019; Presacco et al., 2019; Decruy et al., 2020; Fuglsang et al., 2020). In all these studies, both NH listeners and HI listeners showed a higher neural tracking of the attended speech stream than that of the ignored speech stream. Petersen et al. (2017) reported that adults with a higher degree of hearing loss showed a higher neural tracking of the ignored speech and no change in the attended stream, suggesting that they experience more difficulties inhibiting irrelevant information. Although Mirkovic et al. (2019) and Presacco et al. (2019) did not report a neural difference between the two populations, Decruy et al. (2020) and Fuglsang et al. (2020) observed, in contrast to Petersen et al. (2017), an enhanced neural tracking in HI listeners for the attended-speech compared to their normal-hearing peers. This enhancement can indicate a compensation mechanism: HI listeners need to compensate for the degraded auditory input and therefore show enhanced neural tracking.

By using a forward modeling approach, we aimed to gain more insight into the aforementioned compensation mechanism. An essential factor to consider is the neurophysiological changes that occur as hearing declines. Previous research suggests that due to hearing loss, the brain changes in structure and functionality (for a review, see: Cardin, 2016). These changes in structure and/or function are covered by the concept of neural plasticity, i.e., the ability of the nervous system to change its activity in response to intrinsic or extrinsic stimuli by reorganizing its structure, functions, or connections as defined by Mateos-Aparicio and Rodríguez-Moreno (2019). We aim to evaluate the functional changes by looking at neural tracking and the latency of the neural responses to continuous speech. Subsequently, we investigate whether these neural responses to speech evoke different activation patterns at the sensor level. If so, this indicates that the underlying neural activity is different and therefore indicates structural changes of the brain.

The difficulties of researching HI listeners are twofold. First, most HI listeners are older, and ageing also has an impact on brain responses (Tremblay et al., 2003; Harkrider et al., 2006; Burkard et al., 2007; Harkrider et al., 2009; Decruy et al., 2019; Presacco et al., 2016). Therefore, it is important to compare HI listeners to age-matched normal-hearing peers. Second, audibility of the stimulus must be taken into account: sound presented at the same intensity can be less audible for HI listeners than for NH listeners.

Previous studies which reported the differences between HI listeners and NH listeners, focused on differences in neural tracking, i.e. reconstruction accuracies or prediction accuracies. Firstly, we studied the effect of hearing loss on prediction accuracy, which should align with previous literature: HI listeners show increased prediction accuracies compared to NH listeners (Fuglsang et al., 2020). Subsequently, we investigated whether the neural responses to continuous speech (e.g., latency and topography) differed between HI listeners and NH listeners. By examining characteristics of the neural responses to continuous speech, we aimed to better understand the compensation mechanism, and its associated functional as structural changes, on which HI listeners rely.

## Materials and Methods

### Participants

We used a dataset containing EEG of 14 HI listeners (8 female; average age ±std = 58±20) with sensorineural hearing loss and 14 aged-matched normal-hearing peers (13 female; average age ±std = 51±15). The data were collected in a previous study by Decruy et al. (2020) (medical ethics committee of the University Hospital of Leuven approved the experiment (S57102); all participants signed an informed consent form). Inclusion criteria were: (1) having Dutch as a mother tongue, (2) having symmetrical hearing (i.e., determined based upon the criteria derived from the AMCLASS algorithm of Margolis and Saly (2008)) and (3) absence of medical conditions and learning disorders. A cognitive screening, the Montreal Cognitive Assessment (Nasreddine, 2004), was performed for to ensure the absence of cognitive impairment. Hearing thresholds were determined using pure tone audiometry (125 to 8000 Hz). Normal hearing was defined for all participants where the hearing threshold did not exceed 30 dB HL for frequencies 125 to 4000 Hz (as defined by Decruy et al. (2019); average of hearing thresholds within this frequency range in the stimulated ear is denoted as the pure-tone average (PTA)). The hearing thresholds and PTA are shown in Figure 1 (NH listeners: average PTA ± std= 13.27 ± 5.60 dB HL, HI listeners: average PTA ± std= 44.46 ± 10.54 dB HL).

**Figure 1:**
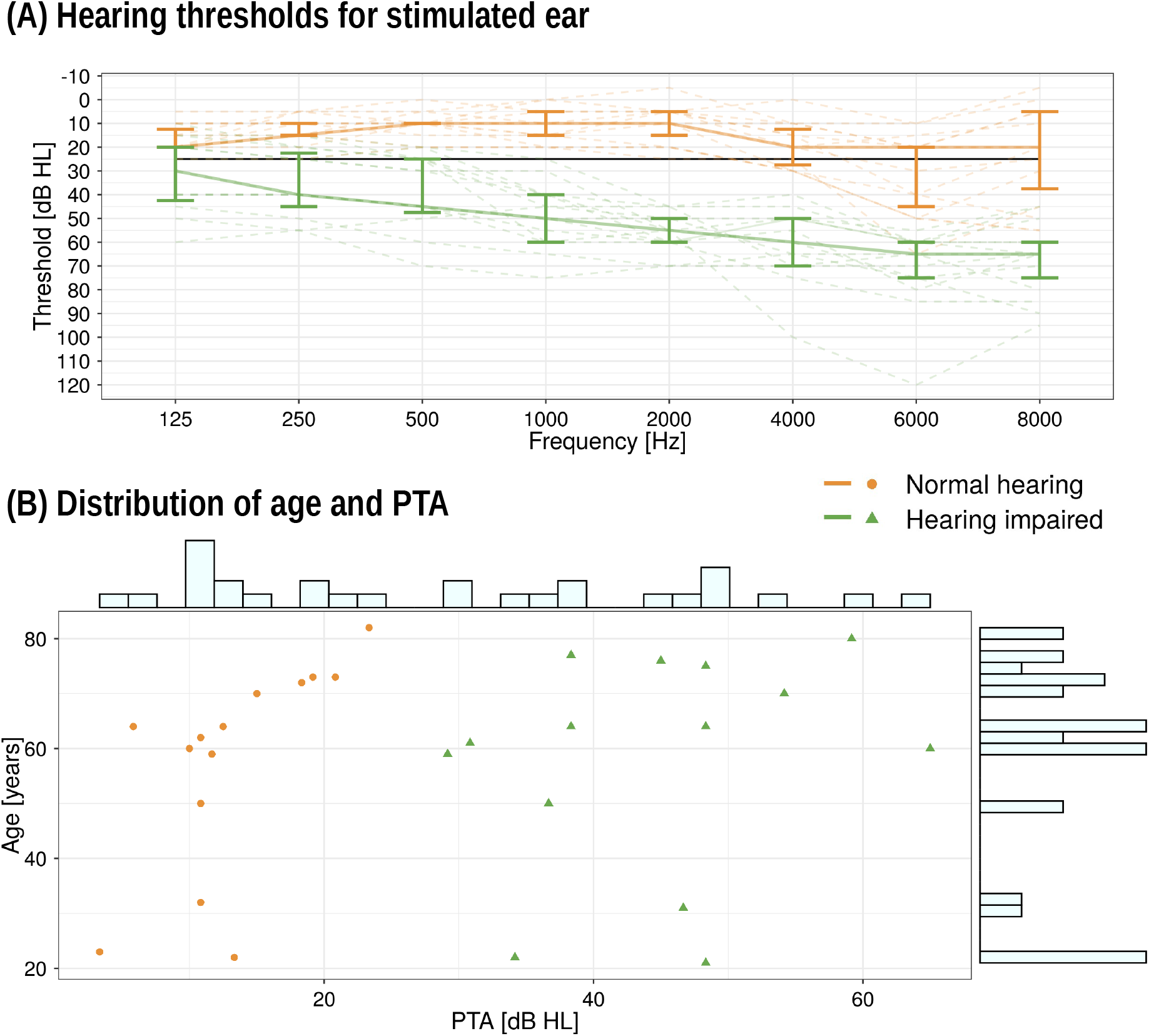
Participant details. Panel A: The hearing thresholds for the stimulated ear. The dashed lines represent the individual hearing thresholds. The full orange or green line annotates the median value for respectively NH listeners and HI listeners. The error bars indicate the range between the 25th and 75th quantile. Panel B: PTA as a function of age for NH listeners (dot, orange) and HI listeners (triangle green). The histograms across the horizontal and vertical axes show the distribution of PTA and age across the participants.

### Experimental Procedures

#### Behavioural Experiment: Flemish Matrix sentence test

The Matrix sentence test was performed to determine the participant’s Speech Reception Threshold (SRT) in speech weighted noise (SWN). These Matrix sentences have a standard grammatical structure, consisting of a name, a verb, a numeral, a colour and an object (Luts et al., 2014). The SRT represents the signal-to-noise ratio (SNR) at which 50% of the presented words are recalled correctly. These sentences were presented at a fixed intensity for the NH listeners (55 dB SPL; A-weighted) and an adjusted intensity for the HI listeners to assure a comfortable level and good audibility (more details in the section Stimuli presentation). To determine the SRT, we used an adaptive procedure that adjusted the background noise level based on the correctly recalled words of the Matrix sentence. We opted to adjust the noise level rather than the speech level to assure that the participant’s speech intelligibility was affected by the noise rather than a decrease in audibility.

#### EEG Experiment

##### Data acquisition

A BioSemi ActiveTwo system (Amsterdam, Netherlands) was used to measure EEG signals during stimuli presentation. This system uses 64 Ag/AgCl electrodes placed according to the 10-20 system (Oostenveld and Praamstra, 2001). The EEG signals were measured with a sampling frequency of 8192 Hz. All recordings were carried out in a soundproof booth with Faraday cage at ExpORL (Dept. Neurosciences, KU Leuven).

##### Stimuli presentation

The speech stimuli were presented monaurally through ER-3A insert phones (Etymotic Research Inc, IL, USA) using the software platform APEX (Dept. Neurosciences, KU Leuven) (Francart et al., 2008). The stimuli were presented to the right ear unless the participant preferred the left ear (n = 3; 1 NH; 2 HI; ear preference was determined with a Flemish, modified version of the laterality preference inventory of Coren (1993)). All stimuli were set to the same root mean square level and were calibrated.

For NH listeners, the speech stimuli’ intensity was fixed at 55 dB SPL (A-weighted). To ensure audible stimuli for HI listeners, the stimuli were linearly amplified based on the participant’s hearing thresholds according to the National Acoustics Laboratory-Revised Profound (NAL) algorithm (Byrne et al., 2001). To ensure a comfortable level, the overall level was adjusted on a subject-specific basis in addition to the linear amplification so that the stimulus was minimally effortful and comfortable to listen to (mean±std = 57.9 ±6.4 dB SPL; A-weighted). Decruy et al. (2020) determined the comfortable loudness level by varying the overall intensity of a story in quiet until participants indicated on a scale that it was comfortable, intelligible, and minimally effortful to understand. The individual presentation levels are reported by Decruy et al. (2020). In the continuation of this manuscript, we use the term *sound intensity* to refer to the SPL levels.

In summary, we compensated for the hearing loss by two types of amplification: (1) we provided frequency-specific amplification of the stimulus based upon their hearing threshold, and (2) we adjusted the overall sound intensity to a comfortable loudness. Without this amplification, listeners with hearing loss perceive a degraded version of the speech and do not have access to the high-frequency information which is crucial for speech intelligibility. Hence, they might allocate more effort to listen the stimulus. Listening to degraded speech (e.g. Mirkovic et al., 2019; Verschueren et al., 2021; Kraus et al., 2020) or spending more effort to listen to the stimulus (e.g. Dimitrijevic et al., 2019) affects the neural responses to continuous speech. By providing amplification, we aimed to make the peripheral activation levels as similar as possible in both groups, which motivates our choice to model the neural responses to the non-amplified speech features for HI listeners.

During the EEG recording, 2 Dutch stories were presented: (1) “Milan”, a 12-minute long story narrated by Stijn Vranken (male) presented in quiet and (2) “De Wilde Zwanen” narrated by Katrien Devos (female) presented in 5 different levels of background speech-weighted noise (each lasted around 2 minutes; SNR conditions were shuffled randomly across the 5 different parts). The duration of silences was limited to 200 ms.

The levels of background noise for the second story depended on the participant’s speech-in-noise performance. As the SRT depends on the presented speech material, we used the self-assessed Békesy procedure to derive the SRT of the story (Decruy et al., 2019). This procedure uses the relation between the subjective rating of the Matrix sentences and the SRT of the Matrix sentences to estimate the story SRT based upon its subjective rating (Decruy et al., 2018, 2019). The noise conditions were calculated on the participant’s story adjusted SRT, namely: SRT - 3 dB, SRT, SRT + 3 dB, SRT + 6 dB and a condition without noise, which approximate speech understanding levels of 20%, 50%, 80%, 95% and 100%. This way the level of speech understanding was kept constant rather than the amount of background noise across the different participants. After each condition, the participant was asked to rate how much they understood on a scale from 0 to 100% to obtain the participant’s subjective rating of speech understanding (Decruy et al., 2018, 2019) (question: *What percentage of the words did you understand correctly?*; subjective ratings are visualized in supplementary Figure S.1).

### Signal Processing

#### Processing of the EEG signals

The EEG recording with a sampling frequency of 8192 Hz was downsampled to 256 Hz to decrease processing time. To remove artefacts of eye blinks, we applied multi-channel Wiener filtering to the EEG data to remove artefacts of eye blinks (Somers et al., 2018). Then we referenced the EEG data to the common-average and filtered the data between 0.5 and 25 Hz using a zero-phase Chebyshev filter (Type II with an attenuation of 80 dB at 10% outside the passband; acausal Chebyshev filtering returns similar outputs as causal Least Squares filtering (order = 500, passband weight = 100, stopband weight = 1, stopband frequency at 10% outside the passband frequency), i.e., similar accuracies (Pearson r > 0.999, p < 0.0001), peak latencies (Pearson r > 0.98, p < 0.0001) and TRF patterns (Pearson r > 0.998)). Additional downsampling to 128 Hz was performed.

#### Extraction of the speech features

In this study, we used 2 speech features: spectrogram and acoustical onsets. Both speech features are continuous features which represent the acoustical properties of the speech stimulus.

To create the spectrogram, the speech stimulus (without amplification) was low-pass filtered below 4000 Hz (zero-phase low-pass FIR filter with a hamming window of 159 samples) because the ER-3A insert phones also low-pass filter at this frequency. A spectrogram representation was obtained using the Gammatone Filterbank Toolkit 1.0 (Heeris, 2014) (centre frequencies between 70 and 4000 Hz with 256 filter channels and an integration window of 0.01 second). This toolkit calculates a spectrogram representation based on a series of gammatone filters inspired by the structure of the human auditory system (Slaney, 1998). The resulting 256 filter outputs were averaged into 8 frequency bands (each containing 32 outputs). Additionally, each frequency band was downsampled to the same sampling frequency as the processed EEG, namely 128 Hz. The NAL filtering introduced a delay of 5.333 ms, which was compensated for by padding the non-delayed speech representation with zeros for the delay’s duration at the stimulus’s beginning (filter delay is constant across frequency). The acoustical onsets were calculated as a half-wave rectification, i.e., negative values were set to zero, of the spectrogram’s derivative.

#### Prediction accuracies, temporal response function & peak picking method

In this study, we focused on a linear forward modeling approach that predicts the EEG based on a linear combination of speech features of the presented speech. This forward modeling approach results in 2 outcomes: (a) a TRF and (b) a prediction accuracy. (a) A TRF is a linear approximation of the brain’s impulse response. It is a signal over time that describes how the brain responds to the speech features. (b) TRFs can be used to predict the EEG by convolving it with the speech features. The predicted EEG is then correlated with the actual EEG to obtain a prediction accuracy. Prediction accuracy is considered a measure of neural tracking: the higher the prediction accuracy, the better the brain tracks the stimulus. A schematic overview of this forward modeling approach is given in Figure 2.

**Figure 2:**
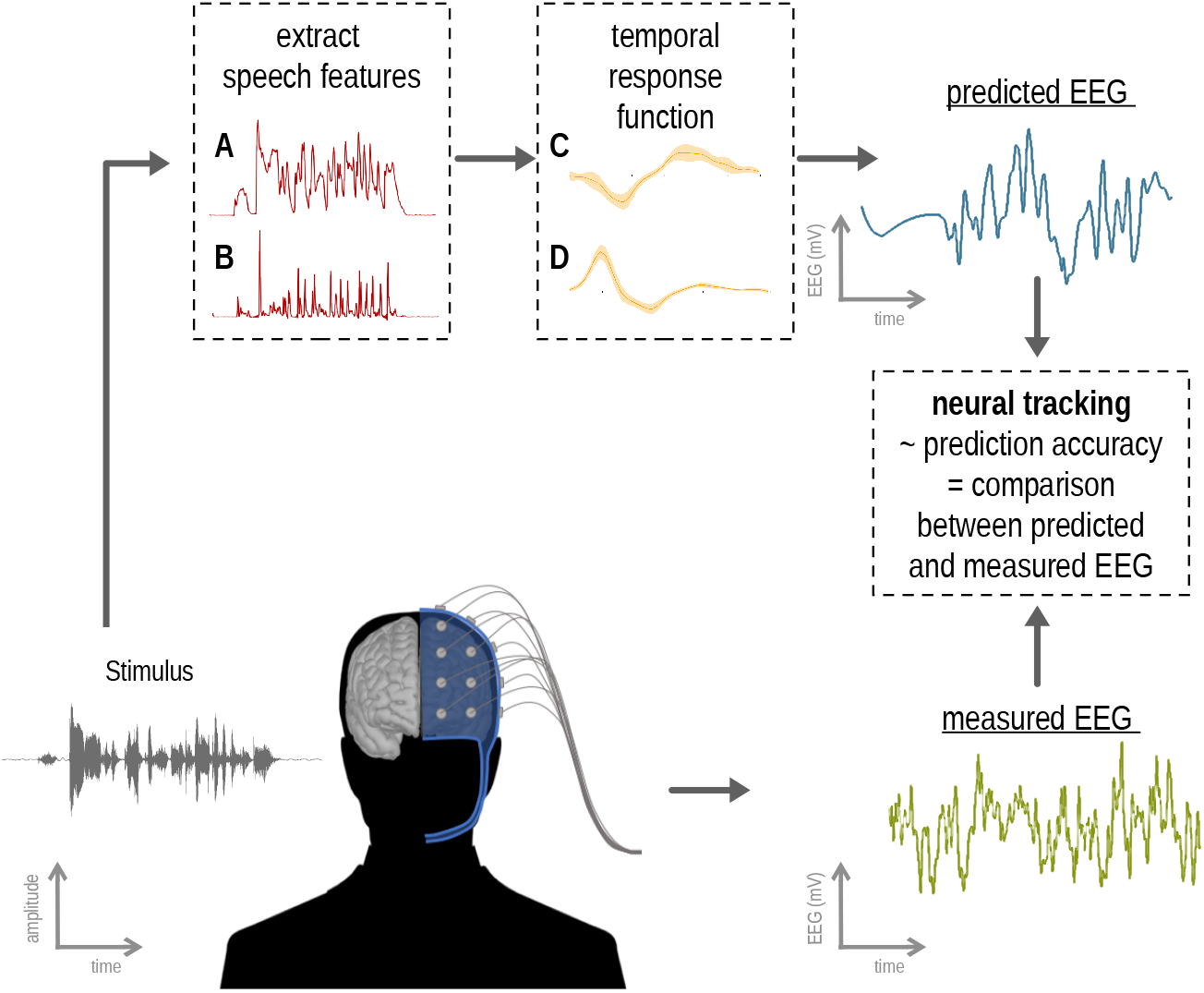
Schematic overview of the forward modeling approach. A participant listens to a stimulus (whether or not presented in noise). From the clean stimulus (i.e., without the noise), we extracted two speech features: the spectrogram (A) and the acoustic onsets (B). For visualization purposes, only one of the eight frequency bands is visualized for these features. The forward modeling approach results in a TRF estimate for each EEG channel and each band of both speech features: one TRF for each channel and each spectrogram band (C) or acoustic onsets band (D). Subsequently, these TRFs are convolved with the stimulus features to predict the EEG signals (for visualization purposes, only one EEG channel is visualized). Neural tracking is then calculated as the correlation between the predicted and measured EEG signal (i.e., a prediction accuracy for each EEG channel).

(a) To estimate TRFs, we used the Eelbrain toolbox (Brodbeck, 2020). The toolbox estimates TRFs using the boosting algorithm by David et al. (2007) (using a fixed step size of 0.005 after Euclidian normalization of the predictor and EEG data; early stopping based on the ℓ_2_-norm by minimizing the Euclidian distance between the actual and predicted EEG data on the validation partition; kernel basis of 50 ms; parameters were kept constant across all participants). We used 4-fold cross-validation (4 equally long folds; 3 folds used for training, 1 for validation) and an integration window between 0 and 700 ms. The estimated TRFs, averaged across folds and frequency bands, were used to determine the peak latencies.

(b) To calculate the prediction accuracy, the TRF is applied to left-out EEG to allow a fair comparison between models with a different number of speech features. We used the boosting algorithm with a testing fold. This implies a 4-fold cross-validation with 2 folds for training, 1 fold for validation and 1 fold for testing, which is left-out during training and validation. Each estimated TRF was used to predict the EEG of the left-out testing fold. The predicted EEG of all left-out segments are correlated, using Pearson correlation, with the actual EEG to obtain a prediction accuracy per EEG-electrode. The prediction accuracies were averaged acros sEEG-electrodes and denoted as neural tracking. Similarly, as Decruy et al. (2020), we calculated neural tracking of the second story, presented in different level of background noise, using the TRFs estimated on the story in quiet.

In summary, to obtain the neural tracking, i.e., prediction accuracy averaged across channels, for the story presented in silence (~13 min data), we used the forward model estimated with testing partition to obtain an unbiased estimate of neural tracking. For the different story parts presented in noise (each ~2 min data), we did not have enough data to estimate a forward model with a testing partition. Therefore, we used the estimated TRF without testing partition of the story in silence to calculate the neural tracking at the different noise levels.

To investigate the spatial and temporal characteristics of the neural responses, we examined the estimated TRF without a testing partition. This way, we used all the data to estimate the TRF, which leads to a better characterization of neural responses to the two speech features.

We did not observe a relation between hearing status and the latencies for the different frequency bands (no significant differences except for amplitude differences, similar to (Brodbeck et al., 2018)). Moreover, we observed that the peak latency for HI listeners is delayed for each frequency band compared to NH listeners. However, for some frequency bands, the pattern of the TRF is less prominent, which is also observed in other studies that investigate the neural response to the spectrogram representation (Di Liberto et al., 2015). As we did not observe an interesting pattern across frequency bands, the TRFs were averaged across the different frequency bands.

From the TRF, we aimed to identify the amplitude and latency of 3 peaks: P1, N1 and P2. As the EEG data contains 64 different channels, 64 different TRFs were estimated, which made peak picking more complex. To investigate the neural changes associated with the development of a hearing loss, we evaluated the neural responses to speech in two ways: by investigating the topographies and the neural response latency. We expected to see differences in topographies between NH listeners and HI listeners, hence, setting one channel selection to determine the peak latency can bias the result. Therefore, we applied principal component analysis (PCA), a dimensionality reduction method: this method allows identifying the relevant channels on a subject-specific basis. The PCA-method results in (a) signals in component space and (b) corresponding spatial filters which describe the linear combinations of EEG channels to obtain these components. In our analysis, the first component was used. Adding more components up to 4 did not change the findings of this study. In addition to the time course of the component, we also investigated the corresponding spatial filter. As the sign of this spatial filter is arbitrary, we forced the average of occipital and parietal channels (P9, P7, PO7, O1, Oz, O2, PO8, P8, Iz, P10) to be negative by multiplying the spatial filter with −1 when needed. This way, we assured that the PCA component had the same polarity across all participants. The PCA-method was applied to the data per story for each participant.

To identify the different peaks, we performed a z-score normalization of the TRF in component space and determined the maximal or minimal amplitude for positive and negative peaks in different time regions (P1: 30 to 110 ms, N1: 70 to 210 ms, P2: 110 to 270 ms), respectively. The overlap of these time regions is not an issue as we identified either the maximal or minimal amplitude to determine the peak latency of a positive or negative peak, respectively. To only identify prominent peaks, a peak was discarded from the analysis if the amplitude of the normalized TRF was smaller than the threshold of 1.

#### Statistical analysis

We used the R software package (version 3.6.3) (R Core Team, 2020) and the Buildmer toolbox, which allows identifying the best linear mixed model (LMM) or linear model (LM) given a series of predictors and all their possible interactions based on the likelihood-ratio test (Voeten, 2020). Depending on the analysis, we used the following predictors: (a) hearing status (NH or HI) or the PTA depending on whether we were interested in the group effect or the effect of the degree of hearing loss, (b) age and (c) peak type (4 levels: P1 and N1 for acoustic onsets, N1 and P2 for spectrogram). To observe an effect of model choice on prediction accuracy, we also included the predictor (d) model type (Spectrogram, Acoustic onsets, Acoustic onsets + Spectrogram) in the statistical analysis. The analysis over different noise conditions also included the predictor (e) speech understanding. For the analyses with regard to peak latency, all continuous predictors were z-scored to minimize effects due to differences in scale. A matching factor indicated the participants belonging to the same age-matched pair. We included a nested random effect: participant nested inside match, as each match contained a pair of participants, and each participant had multiple dependent observations. The models’ assumptions were checked with a visual inspection of the residual plots to assure homoscedasticity and normality. The models’ outcomes were reported with the regression coefficient (*β*) with standard error (SE), t-ratio and p-value per fixed effect. If significant interaction effects were found or if we aimed to identify differences between levels of a factor, additional Holm-adjusted post-hoc tests were performed by looking at the estimated marginal means, linear trends or pairwise comparisons of these estimates, implemented by the Emmeans toolbox (Lenth, 2020). In some instances, to gain more insight into interaction terms, we estimated one predictor’s marginal mean or trend given another continuous predictor. In such cases, we made the latter z-scored predictor discrete to the levels −1 and 1. All post-hoc tests were corrected for multiple comparisons by applying the Holm–Bonferroni method. A significance level of *α* = 0.05 was used.

To compare differences in spatial filters or topographies of the peaks between the 2 groups, we used a related cluster-based permutation test proposed by Maris and Oostenveld (2007) to determine whether the topography differs between NH listeners and HI listeners, using the Eelbrain implementation (Brodbeck, 2020). For these related cluster-based permutation tests the age-matching was preserved. For instance to test whether the topography differed between HI listeners and NH listeners, only the peak topographies of the age-matched participants were considered if both participants showed a prominent peak. A significance level of *α* = 0.05 was used.

## Results

First, we evaluated the effect of hearing loss on the prediction accuracy averaged across channels, further denoted as *neural tracking*. Second, we evaluated the effect of hearing loss on the pattern of the TRF. More specifically, we quantified the effect of hearing loss on the neural response peak latencies and their corresponding peak topographies.

### Neural differences when listening to speech in quiet

#### Differences in neural tracking

We identified the speech feature(s) that resulted in the highest neural tracking: acoustic onsets, spectrogram or a combination of both speech features. As shown in Figure 3 and verified by the statistical analysis, the highest neural tracking was obtained with a combination of both speech features (analysis using LMM: Table 1). Additionally, HI listeners showed higher neural tracking compared to the group of NH listeners (on average 0.012 higher; SE = 0.0054, df = 26, t-ratio = 2.2715, p = 0.0316). Age did not have a significant effect on the neural tracking of speech. A Holm-adjusted pairwise comparison confirmed that the highest neural tracking was obtained with a combination of both speech features which was higher compared to the model using just the acoustic onsets (on average the prediction accuracy of the combined model is 0.003 higher; SE = 0.000651; df = 54, t-ratio = 4.521, p = 0.0001) and higher compared to the model using the spectrogram variable (on average the prediction accuracy of the combined model is 0.005 higher; SE = 0.000651; df = 54; t-ratio = 7.230; p < 0.0001). Therefore, in the continuation of this study, we investigated the corresponding TRFs of the model which combines acoustic onsets with the spectrogram.

**Figure 3:**
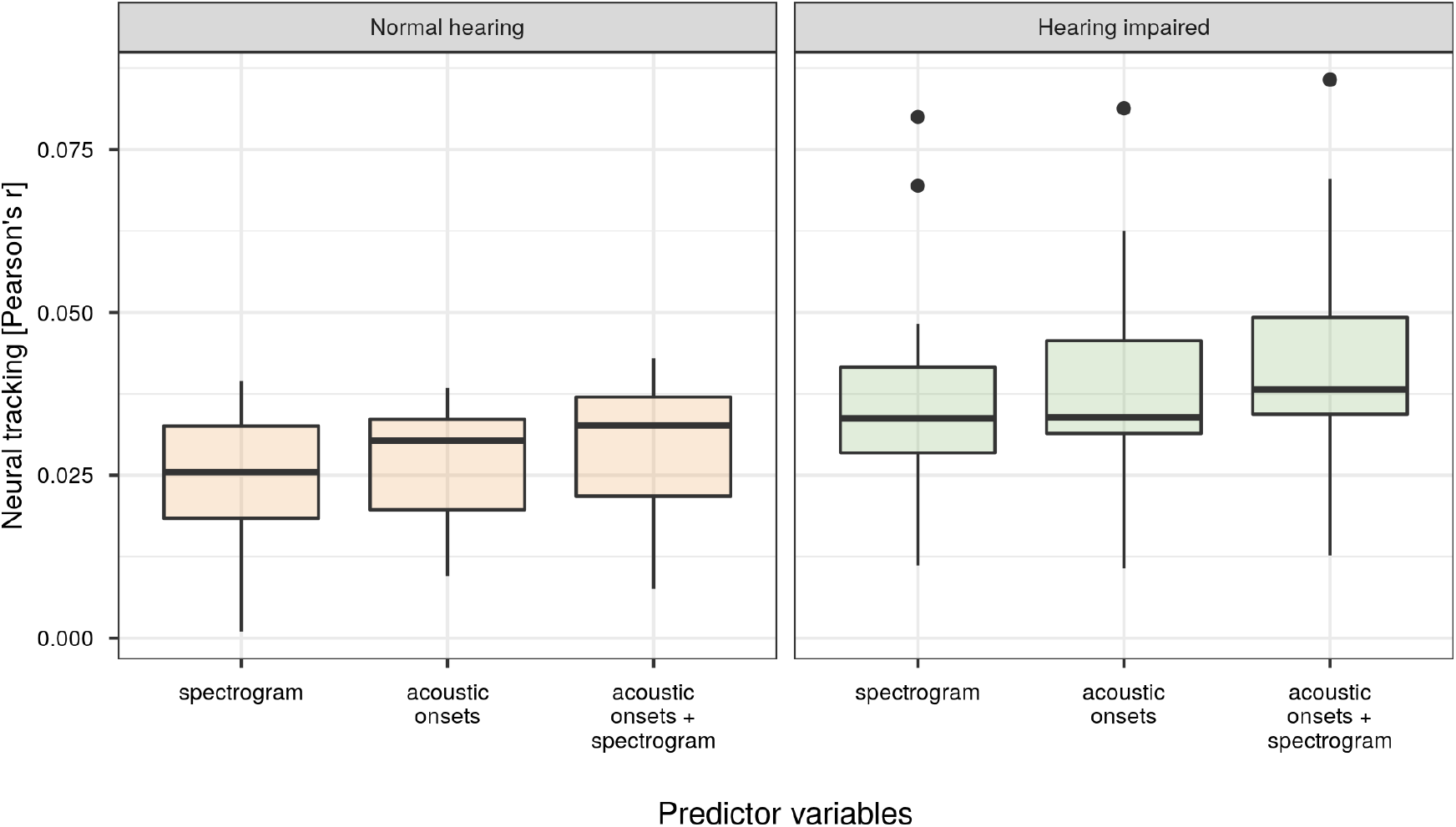
Neural tracking (Pearson’s r) as a function of different combinations of speech features (‘spectrogram’, ‘acoustic onsets’ and ‘acoustic onsets + spectrogram’, respectively) for both NH listeners (left; orange) and HI listeners (right; green).

**Table 1:** Linear mixed model: the effect of hearing status and model type on neural tracking. Estimates of the regression coefficients (*β*), standard errors (SE), degrees of freedom (df), t-Ratios and p-values are reported per fixed effect term. Participant nested in match was included as a random effect. Formula: neural tracking ~ 1 + hearing status + model type + (1 | match/participant)

| fixed effect term | $\beta$ | SE | df | t-Ratio | p-value |
| --- | --- | --- | --- | --- | --- |
| Intercept (for NH / Spectrogram) | 0.0252 | 0.0039 | 26.4961 | 6.5042 | p < 0.001 |
| Hearing status: HL | 0.0124 | 0.0054 | 26 | 2.2715 | p = 0.0316 |
| Model type = Acoustic onsets | 0.0018 | 7e-04 | 54 | 2.7088 | p = 0.0090 |
| Model type = Acoustic onsets + spectrogram | 0.0047 | 7e-04 | 54 | 7.2301 | p < 0.001 |

### Differences in neural response latencies

#### Difference between NH listeners and HI listeners

In Figure 4.A, we visualized the normalized first PCA component of the TRFs for both groups and speech features. The TRFs of HI listeners show delayed neural responses to speech compared to those of NH listeners. Additionally, the average TRF for each speech feature show 2 prominent peaks: P1-peak of acoustic onsets (*P*1_*AO*_) and N1-peak of acoustic onsets (*N*1_*AO*_) for the acoustic onsets, N1-peak of spectrogram (*N*1_*S*_) and P2-peak of spectrogram (*P*2_*S*_) for the spectrogram. Subsequently, we extracted the peak latency of each peak in these TRFs and analyzed whether these peak latencies depended on the considered peak (*P*1_*AO*_, *N*1_*AO*_, *N*1_*S*_, *P*2_*S*_; also referred to as peak type), hearing status (NH listeners or HI listeners) and z-scored age. Using the Buildmer toolbox, we identified that the best LMM to predict the peak latency included main effects of peak type, hearing status and age, and an interaction term between age and peak type (Table 2).

**Figure 4:**
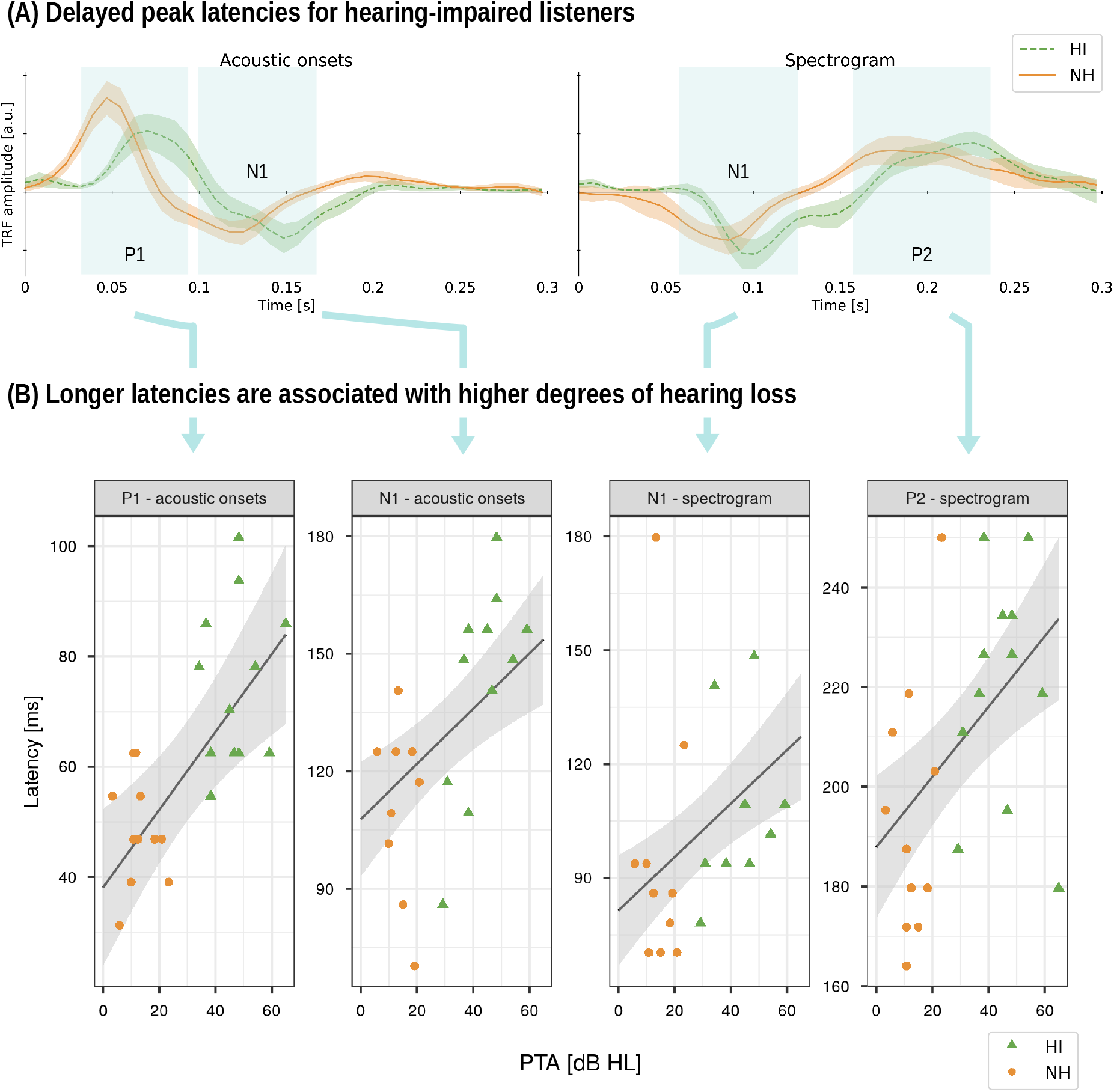
An overview of the neural responses of HI listeners (HI; striped line; green, triangle) and NH listeners (NH; orange, dot). Panel A: normalized first PCA component of the TRFs when listening to a story in quiet for both speech features evaluated for both HI listeners and NH listeners. The thick line represents the average TRF over participants. The shaded area indicates the standard error of this average TRF. Panel B: The peak latency in function of the degree of hearing loss (PTA) derived from the neural responses when listening to a story in quiet.

**Table 2:**
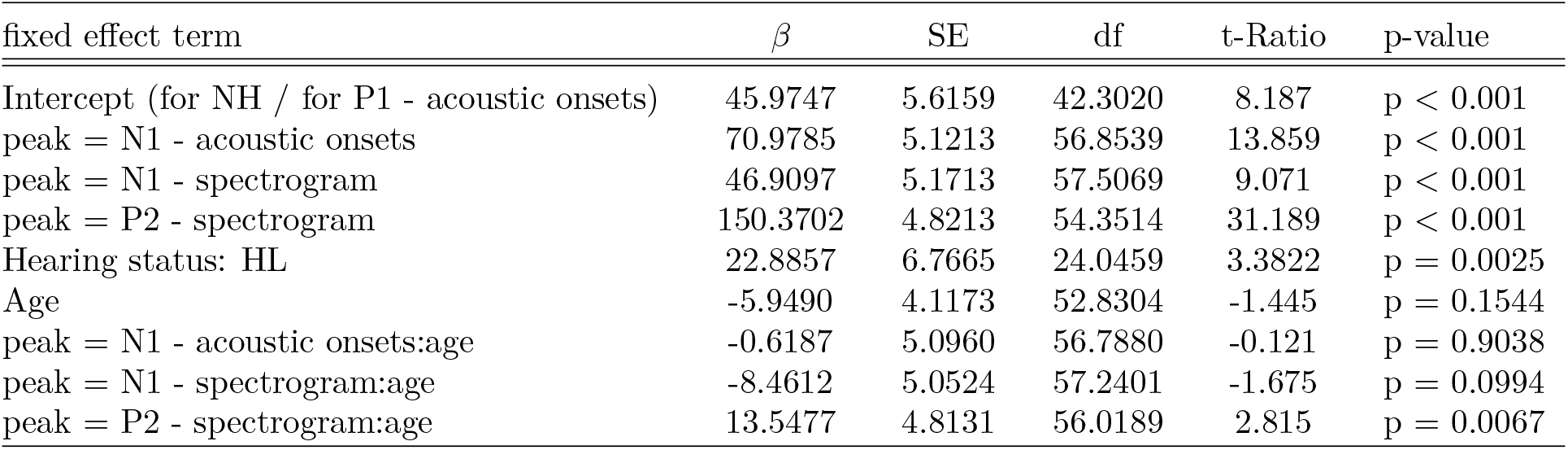
Results of the linear mixed model in order to assess peak type, hearing status and age (z-scored) on the peak latency of *P*1_*AO*_, *N*1_*AO*_, *N*1_*S*_, *P*2_*S*_. Estimates of the regression coefficients (*β*), standard errors (SE), degrees of freedom (df), t-Ratios and p-values are reported per fixed effect term. Participant nested in match was included as a random effect. Formula: latency ~ 1 + peak + hearing status + age + peak:age + (1 | match/participant)

| fixed effect term | $\beta$ | SE | df | t-Ratio | p-value |
| --- | --- | --- | --- | --- | --- |
| Intercept (for NH / for P1 - acoustic onsets) | 45.9747 | 5.6159 | 42.3020 | 8.187 | p < 0.001 |
| peak = N1 - acoustic onsets | 70.9785 | 5.1213 | 56.8539 | 13.859 | p < 0.001 |
| peak = N1 - spectrogram | 46.9097 | 5.1713 | 57.5069 | 9.071 | p < 0.001 |
| peak = P2 - spectrogram | 150.3702 | 4.8213 | 54.3514 | 31.189 | p < 0.001 |
| Hearing status: HL | 22.8857 | 6.7665 | 24.0459 | 3.3822 | p = 0.0025 |
| Age | -5.9490 | 4.1173 | 52.8304 | -1.445 | p = 0.1544 |
| peak = N1 - acoustic onsets:age | -0.6187 | 5.0960 | 56.7880 | -0.121 | p = 0.9038 |
| peak = N1 - spectrogram:age | -8.4612 | 5.0524 | 57.2401 | -1.675 | p = 0.0994 |
| peak = P2 - spectrogram:age | 13.5477 | 4.8131 | 56.0189 | 2.815 | p = 0.0067 |

HI listeners showed later peak latencies (an increase of 23 ms, SE = 6.7665, df = 24.0459, t-ratio = 3.3822, p = 0.0025). Depending on which peak is considered, the effect of age on the peak latency changes. In Table 2, the interaction between age and peak type is shown relative to the age effect for peak *P*1_*AO*_. However, this relative effect does not give insight into the actual trend in the data. Therefore, we performed post-hoc testing to estimate the marginal trends of age for each peak type. This way, we observed that only *N*1_*S*_-latency significantly decreases with increasing age (estimate of marginal trend: −14.41, SE = 4.93, df = 68.8, t-ratio = −2.926, p = 0.0186) while no significant trend was observed for the other peak latencies.

To obtain the TRFs in component space, we applied PCA on the 64 TRFs (one TRF per EEG channel) and used the first PCA component. This method also returns a corresponding spatial filter of this component. We did not observe a significant difference between the spatial filters of HI listeners and NH listeners, which was the case for both speech features (spatial filters are visualized in Figure S.2).

When the peak latencies were determined on the TRF in component space, we extracted the topography of the TRF in sensor space at that latency. Although the spatial filters did not significantly differ between NH listeners and HI listeners, the corresponding peak topographies might. Indeed, HI listeners showed a significantly different topography for *N*1_*S*_ compared to NH listeners (Figure 5). HI listeners showed a more prominent central negativity and a higher occipital positivity which was slightly left-lateralized. However, for the other peak topographies, no significant difference was observed.

**Figure 5:**
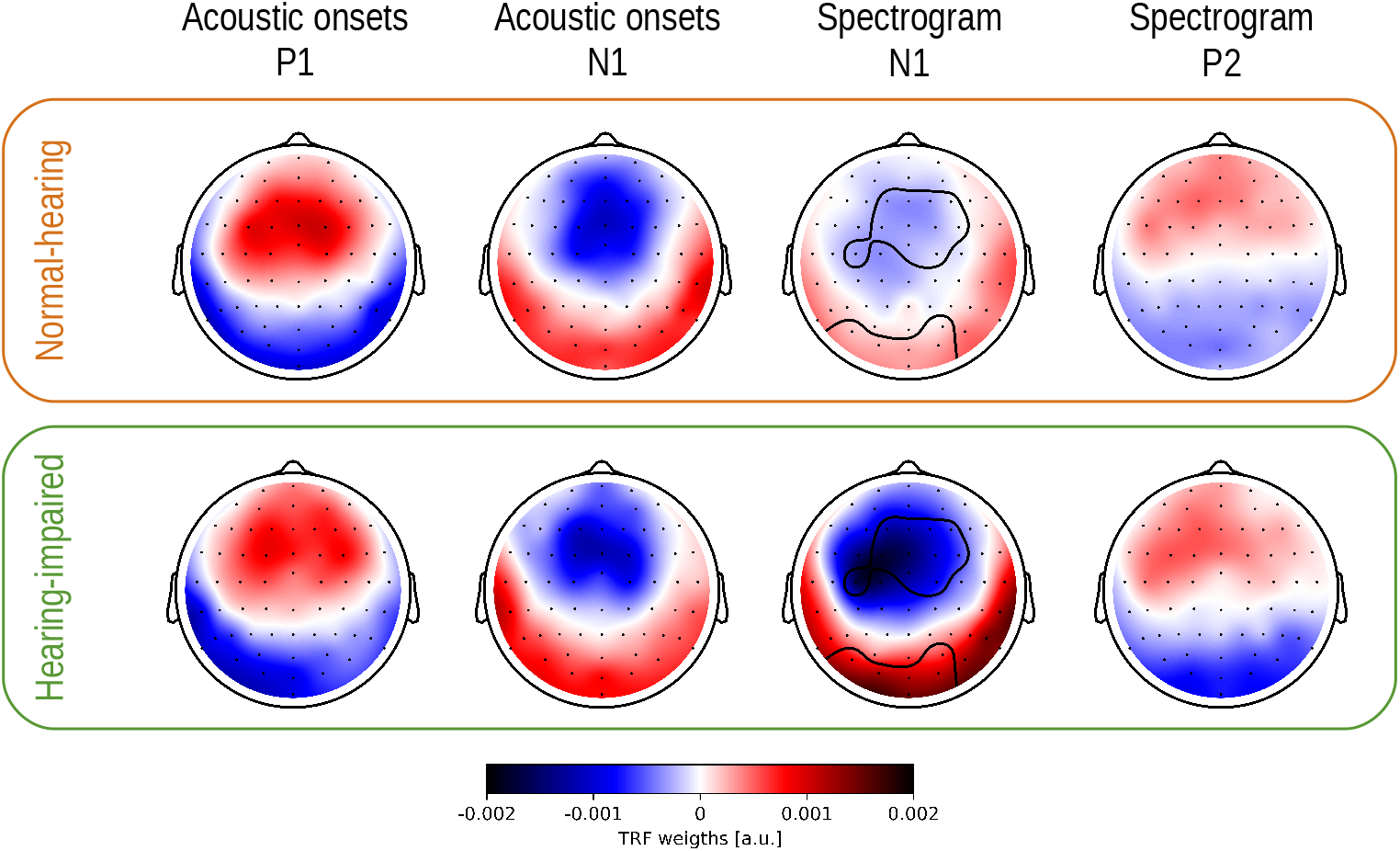
Visualization of the topographies of the peaks in the TRFs in sensor space for both speech features, spectrogram and acoustic onsets, and for NH listeners and HI listeners. The channels which drive the significant difference in the *N*1_*S*_-topography, as identified by the cluster-permutation test, are encircled.

#### Effect of the degree of hearing loss

As significant differences in peak latencies were observed between NH listeners and HI listeners, we hypothesized that a higher degree of hearing loss is associated with increased latency of the peaks. Similarly as above, we identified the LMM which explains the variance in the latency. However, instead of using the factor hearing status, we used the continuous variable describing the degree of hearing loss. We justify this approach because the degree of hearing loss (represented by the PTA) is rather continuously distributed across the participants (Figure 1).

We observed that the peak latency was affected by the peak type, degree of hearing loss (i.e. z-scored PTA value) and age (z-scored) and all its possible interaction terms (Table 3). To investigate how the degree of hearing loss affects the peak latency given the peak type and given the age, we performed a post-hoc test to estimate its marginal trend. The Holm-adjusted estimates of the marginal trend showed that the trend of increasing latency with increasing degree of hearing loss is only significant for older adults for the peak latency of *N*1_*AO*_ (estimate of trend = 21.0, SE = 6.44, df = 61.0, t-ratio = 3.260, p = 0.0146) and *N*1_*S*_ (estimate of this trend = 20.4, SE = 6.38, df = 60.5, t-ratio = 3.192, p = 0.0157). However, this trend did not significantly differ between the different peaks nor between younger and older adults.

**Table 3:** Results of the linear mixed model in order to assess the effects of degree of hearing loss (PTA) and age (z-scored) on the peak latency of *P*1_*AO*_, *N*1_*AO*_, *N*1_*S*_, *P*2_*S*_. Estimates of the regression coefficients (*β*), standard errors (SE), degrees of freedom (df), t-Ratios and p-values are reported per fixed effect term. Participant nested in the matching factor was included as a random nested effect. Formula: latency ~ 1 + peak + PTA + age + peak:age + peak:PTA + peak:PTA:age + (1 | match/participant)

| fixed effect term | $\beta$ | SE | df | t-Ratio | p-value |
| --- | --- | --- | --- | --- | --- |
| Intercept (for P1 - acoustic onsets) | 57.5671 | 4.1993 | 57.7904 | 13.7087 | p < 0.001 |
| peak = N1 - acoustic onsets | 71.513 | 4.9762 | 50.0303 | 14.371 | p < 0.001 |
| peak = N1 - spectrogram | 45.1675 | 5.0515 | 50.412 | 8.9414 | p < 0.001 |
| peak = P2 - spectrogram | 150.0076 | 4.7256 | 47.5163 | 31.7433 | p < 0.001 |
| Degree of hearing loss (PTA) | 14.4906 | 3.9794 | 53.1663 | 3.6414 | p < 0.001 |
| Age | -8.1202 | 3.8447 | 52.9895 | -2.1121 | p = 0.0394 |
| peak = N1 - acoustic onsets:age | -2.2935 | 5.2705 | 53.8308 | -0.4352 | p = 0.6652 |
| peak = N1 - spectrogram:age | -9.1196 | 4.9517 | 51.9372 | -1.8417 | p = 0.0712 |
| peak = P2 - spectrogram:age | 14.7078 | 4.721 | 50.1196 | 3.1154 | p = 0.0030 |
| Degree of hearing loss (PTA):age | 1.1324 | 3.7797 | 52.3099 | 0.2996 | p = 0.7657 |
| peak = N1 - acoustic onsets:PTA | 2.0488 | 5.0542 | 51.1439 | 0.4054 | p = 0.6869 |
| peak = N1 - spectrogram:PTA | -11.919 | 5.3951 | 51.9705 | -2.2092 | p = 0.0316 |
| peak = P2 - spectrogram:PTA | -2.2075 | 4.4718 | 46.5795 | -0.4936 | p = 0.6239 |
| peak = N1 - acoustic onsets:PTA:age | 3.3177 | 5.2119 | 55.1995 | 0.6366 | p = 0.5270 |
| peak = N1 - spectrogram:PTA:age | 16.6746 | 5.7976 | 55.4212 | 2.8761 | p = 0.0057 |
| peak = P2 - spectrogram:PTA:age | -0.5236 | 4.3161 | 47.3511 | -0.1213 | p = 0.9040 |

Figure 1 shows that age is not evenly distributed. Therefore, the age effects in the analysis above might be biased towards three younger age-matched pairs. Hence, we replicated the analysis without these three pairs. Using this subset of the data, the best LMM to explain the peak latency did not contain the three-way interaction between age, degree of hearing loss and peak type, nor the interaction between peak type and degree of hearing loss. However, an interaction between age and peak type was still observed. More specifically, how age affects the peak latency of P1ao is significantly different from the age trend seen for the P2s latency (Table S.1). To gain more insight into this interaction between age and peak type, we performed post hoc tests to estimate the marginal trends of age for each peak type. These Holm-adjusted estimates of the effect of age on the peak latency were not significant for any of the peak latencies. This analysis indicates that the observed age effect is not robust: firstly, it is biased by these three age-matched younger pairs, and secondly, although the age trends for the different peaks differ, the post hoc test showed that the marginal trends do not reach significance. Therefore, this effect was not visualized in Figure 4.

However, it is reassuring to observe that even when taking a subset of the dataset, we observe that the peak latency is affected by the degree of hearing loss (estimate = 12.895, SE = 3.518, df = 17.872, t-ratio = 3.665, p = 0.002). This observed trend was independent of age and peak type.

To verify whether the effect of the degree of hearing loss on the peak latency is robust, we repeated the above analysis using only HI listeners. Although this reduced the statistical power, we observed a significant effect of degree of hearing loss on the *N*1_*AO*_-latency: HI listeners with a more severe hearing loss showed an increased latency (analysis using LM and scaled predictors; Table S.2; estimate = 34.89, SE = 12.20, t-ratio = 2.860, p = 0.0188). We did not observe a significant effect of hearing loss or age on the peak latency for the other peak types.

### Neural differences when speech understanding decreases

#### Differences in neural tracking

The effect of increased neural tracking for HI listeners was robust over different levels of background noise (estimate = 0.0154, SE = 0.0046, df = 25.1566, t-ratio = 3.3594, p = 0.0025); Table 4; Figure 6). Additionally, higher neural tracking was observed with increasing age (estimate = 3e-04, SE = 1e-04, df = 24.9812, t-ratio = 2.2452, p = 0.0339); Table 4; Figure 6) and with increasing speech understanding (estimate = 2e-04, SE = 0, df = 115.3178, t-ratio = 5.6547, p < 0.001; Table 4; Figure 6). No significant interaction effect was observed between hearing status and speech understanding.

**Figure 6:**
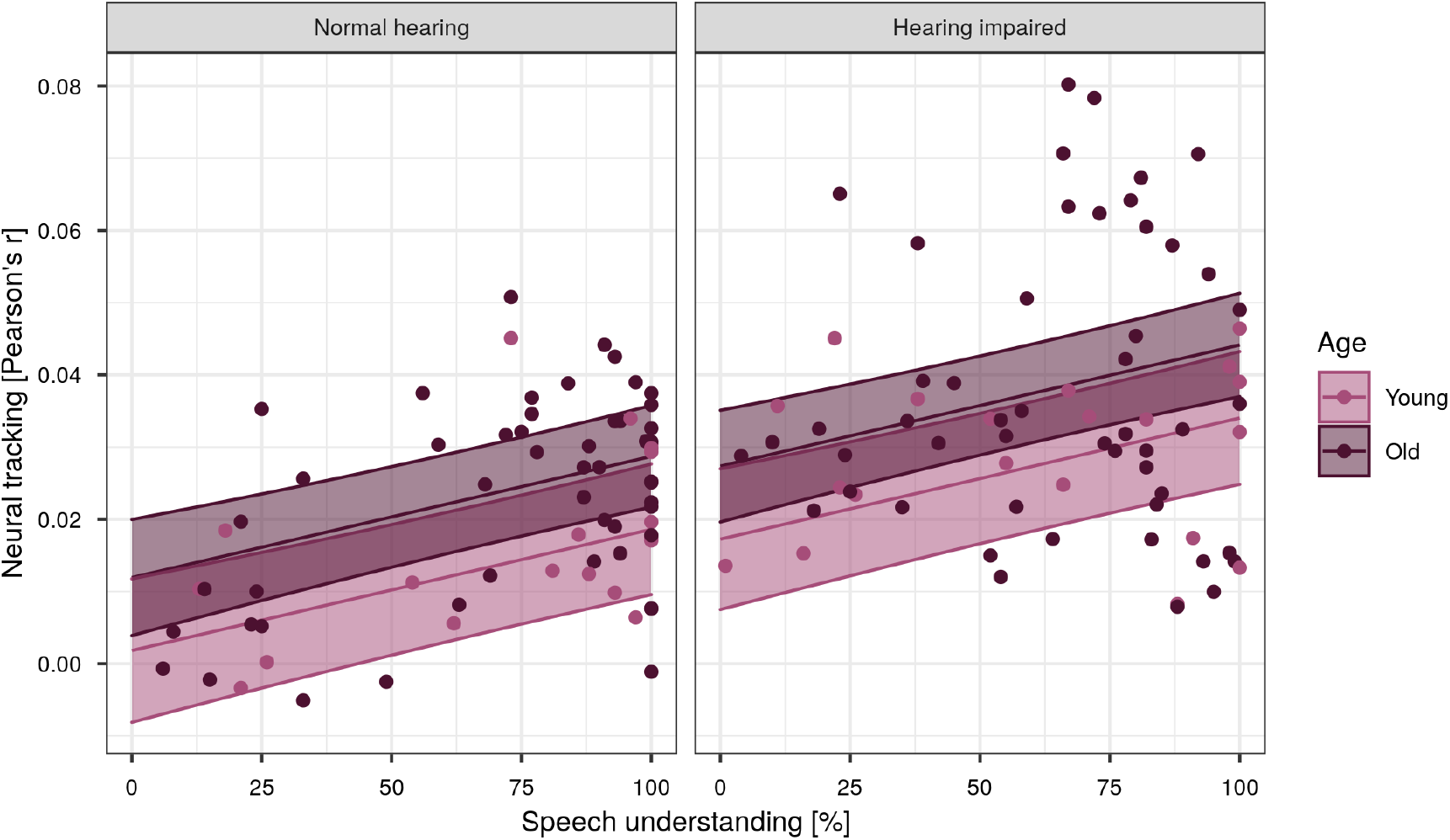
Neural tracking (Pearson’s r) as a function of speech understanding. For visualisation purposes, the effect of age was discretized with 2 levels: the average age of participants younger than 50 years (level young; 31 years; pink) and average age of participants older than 50 years (level old; 68 years; purple) for NH listeners and HI listeners. The level of 50 years was set by visual inspection of the age distribution (Figure 1).

**Table 4:** Linear mixed model: the effect of hearing status and speech understanding on neural tracking. Estimates of the regression coefficients (*β*), standard errors (SE), degrees of freedom (df), t-Ratios and p-values are reported per fixed effect term. Participant nested in match was included as a random effect. Formula: neural tracking ~ 1 + hearing status + age + speech understanding + (1 | match/participant)

| fixed effect term | $\beta$ | SE | df | t-Ratio | p-value |
| --- | --- | --- | --- | --- | --- |
| Intercept (for NH) | -0.0068 | 0.0081 | 29.4306 | -0.8397 | p = 0.4078 |
| Hearing status: HL | 0.0154 | 0.0046 | 25.1566 | 3.3594 | p = 0.0025 |
| Age | 3e-04 | 1e-04 | 24.9812 | 2.2452 | p = 0.0339 |
| Speech understanding | 2e-04 | 0 | 115.3178 | 5.6547 | p < 0.001 |

### Differences in neural response latencies

#### Effect of the degree of hearing loss

Subsequently, we analysed how the neural response latency is affected by speech understanding, age and the degree of hearing loss for the second story presented in different levels of background noise. From the above analysis using speech in silence, we observed that the degree of hearing loss influences the peak latencies. Therefore, we here immediately consider the effect of hearing loss as a continuous variable, i.e. degree of hearing loss, instead of a categorical predictor, i.e. hearing status.

Using Buildmer, we identified the best LMM to explain the second story’s peak latencies, which was presented in different levels of background noise. This model shows that the peak latency of the neural responses depends on the peak type, age and degree of hearing loss (Table 5; Figure 7.B). Moreover, we observed interactions between speech understanding and degree of hearing loss, between peak type and speech understanding, between degree of hearing loss and age, between peak type and age and speech understanding and age. These interactions show that the effect of age and speech understanding on peak latency depended on the peak type. However, by looking at the estimates of these age effects, we observe that these are not consistent across the different peak types. Furthermore, from the analysis using speech in silence, we observed that the age effect is not a robust effect. Indeed, post-hoc tests showed that the marginal trends of age on the peak latency did not show a significant effect for any of the peak types. Therefore, this effect is not visualized in Figure 7.B.

**Figure 7:**
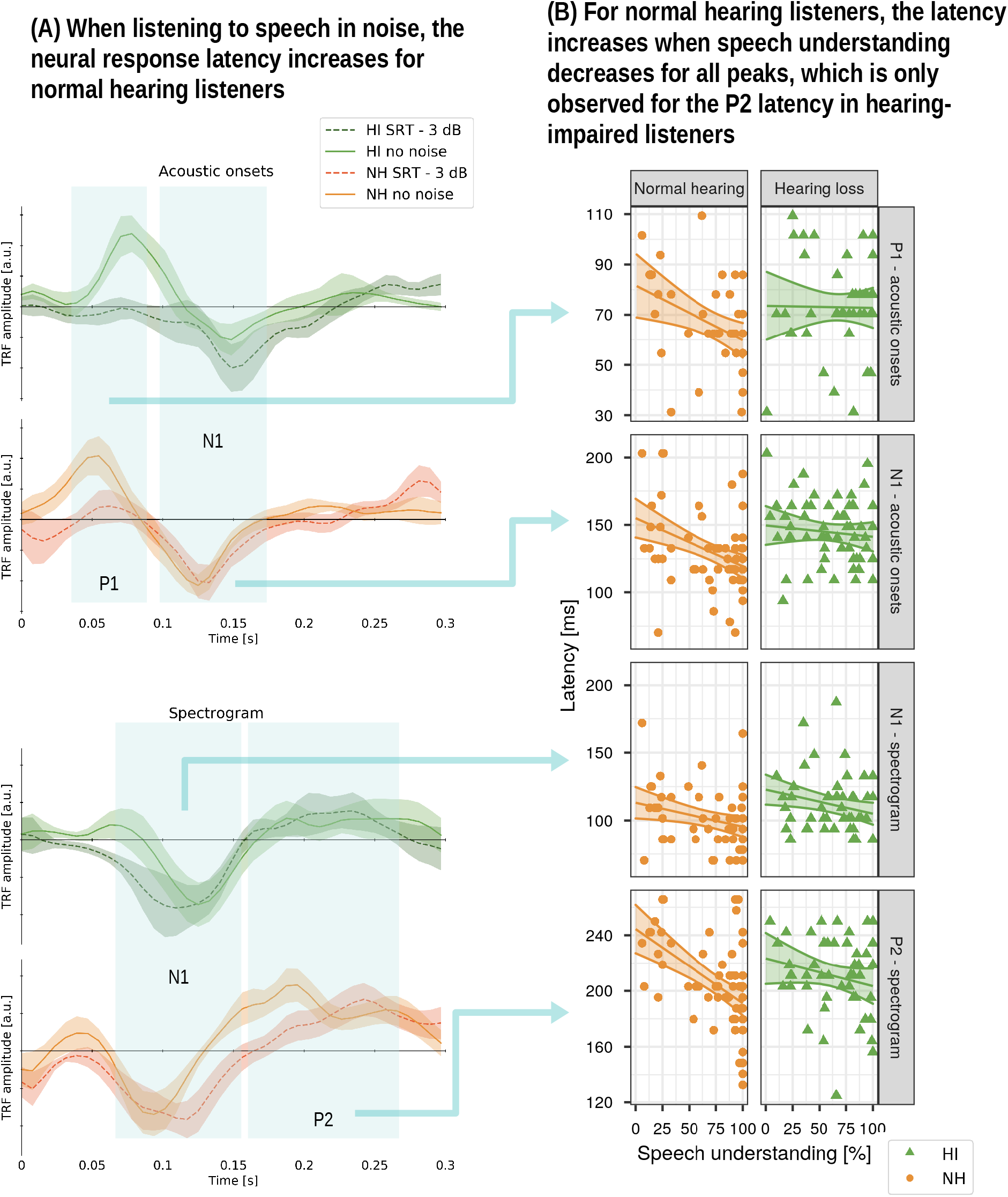
An overview of the neural responses of HI listeners (HI; green, triangle) and NH listeners (NH; orange, dot) when listening to speech in silence and speech presented in noise. Panel A: normalized first PCA component of the TRFs when listening to a story part in quiet (full line) and a story part in noise (striped line) for both speech features evaluated for both HI listeners and NH listeners for visualization purpose only the lowest SNR is visualized, namely SRT - 3 dB). The shaded area indicates the standard error of the average TRF which is represented by the full line. Panel B: The corresponding peak latencies in function of speech understanding. The effect of degree of hearing loss was made discrete at 2 levels, the average hearing thresholds of all NH listeners (level NH listeners: 13 dB HL) and subjects with hearing loss (level HI listeners: 44 dB HL) and is represented by the regression lines with confidence intervals (shaded area).

**Table 5:** Results of the linear mixed model in order to assess the effects of degree of hearing loss (z-scored), speech understanding (z-scored) and age (z-scored) on the peak latency of *P*1_*AO*_, *N*1_*AO*_, *N*1_*S*_, *P*2_*S*_. Estimates of the regression coefficients (*β*), standard errors (SE), degrees of freedom (df), t-Ratios and p-values are reported per fixed effect term. Participant nested in the matching factor was included as a random nested effect. Formula: latency ~ 1 + peak + SI + degree of hearing loss + speech understanding:degree of hearing loss + peak:speech understanding + age +degree of hearing loss:age + peak:age + speech understanding:age + (1 | match/participant)

| fixed effect term | $\beta$ | SE | df | t-Ratio | p-value |
| --- | --- | --- | --- | --- | --- |
| Intercept (for P1 - acoustic onsets) | 65.523 | 3.149 | 69.524 | 20.808 | p < .001 |
| peak = N1 - acoustic onsets | 71.012 | 3.015 | 384.446 | 23.557 | p < .001 |
| peak = N1 - spectrogram | 39.875 | 3.203 | 385.734 | 12.449 | p < .001 |
| peak = P2 - spectrogram | 144.887 | 3.097 | 385.778 | 46.783 | p < .001 |
| Speech understanding | -2.46 | 2.336 | 388.475 | -1.053 | p = 0.293 |
| Degree of hearing loss (PTA) | 5.489 | 2.324 | 23.74 | 2.362 | p = 0.027 |
| Age | -6.421 | 2.882 | 59.27 | -2.228 | p = 0.03 |
| Speech understanding:Degree of hearing loss (PTA) | 4.441 | 1.071 | 396.822 | 4.148 | p < .001 |
| peak = N1 - acoustic onsets:speech understanding | -4.487 | 2.968 | 381.779 | -1.512 | p = 0.131 |
| peak = N1 - spectrogram:speech understanding | -1.148 | 3.163 | 382.021 | -0.363 | p = 0.717 |
| peak = P2 - spectrogram:speech understanding | -8.703 | 3.066 | 383.334 | -2.839 | p = 0.005 |
| Degree of hearing loss (PTA):Age | 4.797 | 2.232 | 24.283 | 2.149 | p = 0.042 |
| peak = N1 - acoustic onsets:age | 4.906 | 2.832 | 379.405 | 1.732 | p = 0.084 |
| peak = N1 - spectrogram:age | 0.94 | 3.196 | 384.574 | 0.294 | p = 0.769 |
| peak = P2 - spectrogram:age | 13.103 | 2.897 | 384.584 | 4.523 | p < .001 |
| Speech understanding:Age | -2.121 | 1.082 | 390.971 | -1.96 | p = 0.051 |

Interestingly, a significant interaction effect between speech understanding and degree of hearing loss was found (estimate = 4.441, SE = 1.071, df = 396.822, t-ratio = 4.148, p < .001). Since there is no three-way interaction between peak type, speech understanding, and the degree of hearing loss, this indicates that interaction between the degree of hearing loss and speech understanding is similar for all peak types. The effect of speech understanding on the peak latency has a negative estimate, which implies that with increasing speech understanding, the latency decreases. This trend is consistent for all four peak types. The interaction term’s estimate is positive and indicates that as the degree of hearing loss increases, the effect of speech understanding on the peak latency becomes less negative and slightly flattens out, which is also visible in Figure 7.

We performed a post-hoc test to estimate the marginal trend of speech understanding for each peak type and hearing loss to gain more insight into this effect. Indeed, as observed in Figure 7: with increasing speech understanding, latency decreases for NH listeners in all peak types (P1ao: estimate = −6.902, SE = 2.50, df=386, t-ratio = −2.760, p-value = 0.0303; N1ao: estimate = −11.388, SE = 2.14, df = 385, t-ratio = −5.310, p-value < 0.0001; N1s: estimate = −8.050, SE = 2.32, df = 385, t-ratio = −3.469, p-value = 0.0035; P2s: estimate = −15.605, SE = 2.23, df = 384, t-ratio = −6.991, p-value < 0.0001), while this effect for HI listeners is only observed for the P2s-latency (estimate = −6.722, SE = 2.44, df = 393, t-ratio = −2.756, p-value = 0.0303) and not for other peak types.

The effect of the degree of hearing loss on the peak amplitude was not consistent for all peaks and therefore not elaborated upon in this manuscript.

## Discussion

We compared the neural responses to continuous speech of adults with a sensorineural hearing loss with those of age-matched normal-hearing peers. We found that HI listeners show higher neural tracking and increased peak latencies in their neural responses. Across noise conditions, NH listeners showed increased latencies as speech understanding decreased. However, for adults with hearing loss, this increase in latency was not observed.

### Higher neural tracking of speech in hearing-impaired listeners

By evaluating neural tracking, we concluded that (1) higher neural tracking is observed for a combination of the spectrogram and acoustic onsets compared to the speech features individually and (2) HI listeners show enhanced neural tracking compared to normal-hearing peers.

The combination of the two speech features results in higher neural tracking than the individual features alone. Therefore, it suggests that the acoustic onsets describe different neural response characteristics than the spectrogram and vice versa. Following Hamilton et al. (2018) and Brodbeck et al. (2020), we hypothesize that both speech features allow a differentiation between sustained activity, i.e. ongoing sounds, represented by the spectrogram, and transient activity, i.e. onset of the sound, represented by acoustic onsets.

Using a forward instead of a backward modeling approach, we observed enhanced neural tracking in HI listeners. Although the conclusions of a forward and backward model are expected to converge, it is reassuring to observe that the conclusions align even though we used different speech features and a different modeling approach. Accordingly, using the same dataset but a different modeling approach, our results agree with those of Decruy et al. (2020). Also Fuglsang et al. (2020) reported that HI listeners have higher neural tracking than NH listeners of the attended speaker. Nevertheless, Presacco et al. (2019) did not find a difference in neural tracking between the two populations. However, in their study, the populations were not closely age-matched, while ageing is known to increase neural tracking (Presacco et al., 2016; Decruy et al., 2019).

Like previous literature, we also observed that neural tracking decreases with decreasing speech understanding (Vanthornhout et al., 2018; Lesenfants et al., 2019; Decruy et al., 2020). Even when the speech is presented with background noise, HI listeners showed enhanced neural tracking of speech. This suggests evidence for a compensation mechanism: higher neural tracking indicates more neural activity to compensate for the degraded auditory input (Eggermont, 2017; Fuglsang et al.,2020). Although neural tracking of speech was enhanced in HI listeners, the effect of speech understanding on the neural tracking was similar for both populations. We also observed an effect of age: older adults showed higher neural tracking compared to younger adults. Although this converges with the results of ageing studies (Presacco et al., 2016; Decruy et al., 2019), we are hesitant to interpret this effect, as in subsequent analyses, we inferred that the age effects are biased towards the three younger age-matched pairs (Figure 1).

### Hearing-impaired listeners process speech less efficiently

HI listeners showed significantly increased latencies compared to their age-matched normal-hearing peers when they listened to a story presented in quiet (Figure 4.A). Additionally, the delay in neural responses increased with a higher degree of hearing loss (Figure 4.B).

Investigating the CAEP-response to syllables, Campbell and Sharma (2013) and Bidelman et al. (2019b) have reported an increased P2 latency with worse speech perception in noise (QuickSIN scores) but not with the degree of hearing loss. However, in both studies, the speech was presented at the same intensity to both HI listeners and NH listeners. McClannahan et al. (2019) remarked that differences in the audibility of the stimulus might explain the differences in neural response latency: at the same intensity, HI listeners will have a worse perception of the speech compared to NH listeners. Indeed, Verschueren et al. (2021) concluded that when the audibility of the stimulus decreases, the latency of the neural response to continuous speech increases. However, this is only true at intensities where the speech audibility affects the listening effort and speech understanding. For NH listeners, the latency reaches a plateau at a comfortable loudness (intensities of 60 dB or higher). Here, we amplified the stimulus based on the participants’ hearing thresholds and presented the speech at a subject-specific intensity to assure comfortable listening for HI listeners. Since we compared the latency of the neural responses at a comfortable loudness, i.e. when the latency has reached a plateau, we can assume that the increased latencies for HI listeners are due to the impact of the hearing loss rather than the impact of a different speech intensity.

Even though the sound was amplified, HI listeners showed increased latencies. Therefore, we hypothesize that there are some intrinsic differences in neural speech processing between HI listeners and NH listeners.A possible explanation may be that HI listeners process speech less efficiently, as proposed by Bidelman et al. (2019a). Using functional connectivity analysis, Bidelman et al. (2019a) showed that HI listeners have higher eccentricity networks, i.e. brain areas communicate in a chain-like fashion rather than star-like. Higher eccentricity suggests (a) more long-range neural signalling, i.e. more extended neural communication pathways, and (b) less efficient information exchange among different brain regions. (a) More extended neural communication pathways may reflect a form of compensation in which additional brain regions are recruited to understand the degraded auditory input. This is supported by increased frontal activation in HI listeners in the neural responses to simple sounds (Campbell and Sharma, 2013; Bidelman et al., 2019b). Similarly, using continuous speech rather than simple repeated sounds, we showed that HI listeners have a significantly different *N*1_*S*_ peak topography. Such a peak topography reflects the sensor response to the activity of underlying neural sources, which respond in a time-locked fashion to the speech material. Therefore, a significant difference in topography suggests the recruitment of additional or different underlying neural sources (Figure 5). (b) When more or different brain regions are involved to process the speech, it causes longer communication pathways in the brain and therefore decreases the neural speech processing efficiency (Bidelman et al., 2019b). Here, we propose the neural response latency as a marker for the efficiency of neural processing of continuous, natural speech: less efficient speech processing is reflected by increased neural response latency as information exchange is hampered due to more involved brain regions and longer communication pathways.

In summary, even though the sound was amplified, HI listeners rely on a compensation mechanism: more brain regions are involved to understand the speech, which decreases the efficiency of the neural speech processing and increases the neural response latency. The efficiency of neural speech processing decreases as the severity of the hearing loss increases. Therefore, we hypothesize that this compensation mechanism is a gradual effect which occurs as the hearing deteriorates.

When the speech understanding decreases, NH listeners showed a prominent increase in latency, while this was less prominent for HI listeners. In several studies, it has been shown that NH listeners show an increased neural response latency with increasing task demand due to lower stimulus intensity, increasing background noise or stimulus vocoding. This is the case for neural processing of continuous speech (Mirkovic et al., 2019;Verschueren et al., 2021; Kraus et al., 2020) as well as simple sounds (Billings et al., 2015; Van Dun et al., 2016; Maamor and Billings, 2017; McClannahan et al., 2019).

However, we did not observe this increase in latency with increasing task demand in adults with a hearing loss (Figure 7.B). We can integrate these findings with the results of Bidelman et al. (2019a). Although Bidelman et al. (2019b) did not report an increase in P2-latency when the stimulus was noise-degraded, in the same data, Bidelman et al. (2019a) reported that when the stimulus was noise-degraded, NH listeners showed more long-range neural signalling, whereas this was not seen for HI listeners (Bidelman et al., 2019a). Our data support the latter finding: in NH listeners the neural response latency increases as the speech understanding decreases due to increasing level of background noise, while this was less prominent for HI listeners. Following the above reasoning: as the stimulus is noise-degraded, the task demand increases. Therefore, more processing time is required to attend the speech stream and ignore the noise, which decreases the efficiency of neural auditory processing, i.e. the response latency is a marker for the efficiency of neural processing of speech. However, this effect is only observed for NH listeners.

For HI listeners, this is not the case: when background noise increases, processing efficiency does not decrease, i.e. no increase in latency, but remains elevated. This aligns with the finding of Bidelman et al. (2019a): they reported that HI listeners did not show more long-range neural signalling when the stimulus was noise-degraded. Assuming that longer latencies are a marker for less efficient neural processing and thus the number of recruited brain regions: our results suggest that HI listeners recruit already a large number of brain regions to understand speech in quiet. Their neural response latency does not increase with an increasing amount of background noise, indicating that no additional brain regions could be required to compensate for the increased demand.

Our findings explain why Mirkovic et al. (2019) did not find a difference in latency between NH listeners and HI listeners as they presented only two noise conditions. As the noise level increases, the difference in latency between the two populations becomes smaller, which reduces the likelihood of a statistical difference between the two populations.

A caveat of the current study is that the age distribution across the different age-matched pairs is not uniform. Therefore, we are hesitant to interpret the current age effects (or interactions with age) in the current dataset as these might be biased towards the younger-age matched pairs. We suggest that future research should investigate the effect of hearing loss on the neural responses by exploring a dataset with age-matched pairs, which are uniformly distributed across the age range. This way, the effect of age and hearing loss on the neural responses to continuous speech can be disentangled. We have to note, however, that finding young HI listeners is challenging. Moreover, the aetiology of hearing loss should also be taken into account as it might affect how the brain responds to continuous speech.

Finally, we would like to highlight the difference in the trend of neural tracking and neural response latency. As speech understanding decreases, neural tracking decreases for both NH and HI listeners while the neural response latency remains constant (HI) or increases (NH). This difference in trend suggests that both measures capture different underlying neural processes for speech comprehension.

## Conclusion

In this study, we compared the neural responses to continuous speech of adults with a sensorineural hearing loss with those of age-matched normal-hearing peers. HI listeners showed increased peak latencies of their neural responses. Interestingly, the latency increases as the degree of hearing loss increases. Across noise conditions, latency generally increases as the listening conditions become more difficult. However, for HI listeners, this increase in latency is not observed. We here suggest latency as a marker for the efficiency of neural processing to understand continuous, natural speech.

## Acknowledgements

The authors would like to thank all members of the ExpORL ISIFIT team for their weekly guidance. Furthermore, we would like to thank Christian Brodbeck. He helped us with applying the Eelbrain toolbox, which accelerated this research project.

## Contributions

M.G., L.D., J.V., and T.F. designed and performed research. M.G. and L.D. analysed data. M.G., L.D., J.V., and T.F. wrote the paper.

## Data availability statement

The data that support the findings of this study can be made available upon request, in so far as this is in agreement with privacy and ethical regulations.

## Acronyms

*N*1_*S*_: N1-peak of spectrogram. 14, 15, 17, 18, 21, 25, 34
*N*1_*AO*_: N1-peak of acoustic onsets. 14, 15, 17, 18, 21, 34
*P*1_*AO*_: P1-peak of acoustic onsets. 14, 18, 21, 34
*P*2_*S*_: P2-peak of spectrogram. 14, 18, 21, 34
*β*: regression coefficient. 11
CAEP: cortical auditory evoked potentials. 2, 24
EEG: electroencephalography. 2, 4, 6-10
HI: listeners hearing impaired listeners. 2-7, 10, 12-17, 19, 20, 22-26, 34
LM: linear model. 11, 17
LMM: linear mixed model. 11, 12, 15, 20
N1: first negative peak. 2, 10, 11
NH listeners: normal hearing listeners. 2-6, 10, 12-17, 19, 20, 22-26
P1: first positive peak. 2, 10, 11
P2: later positive peak. 2, 10, 11, 24, 25
PCA: principal component analysis. 10, 11
PTA: pure-tone average. 4, 5, 11, 15, 16
SE: standard error. 11
SNR: signal-to-noise ratio. 6, 7
SRT: Speech Reception Threshold. 5-7
SWN: speech weighted noise. 5
TRF: temporal response function. 3, 8-12, 14-17, 22 [title=Abbreviations]

## Supplementary Material

**Participant Details: subjective rating for each SNR condition**

**Figure S.1:**
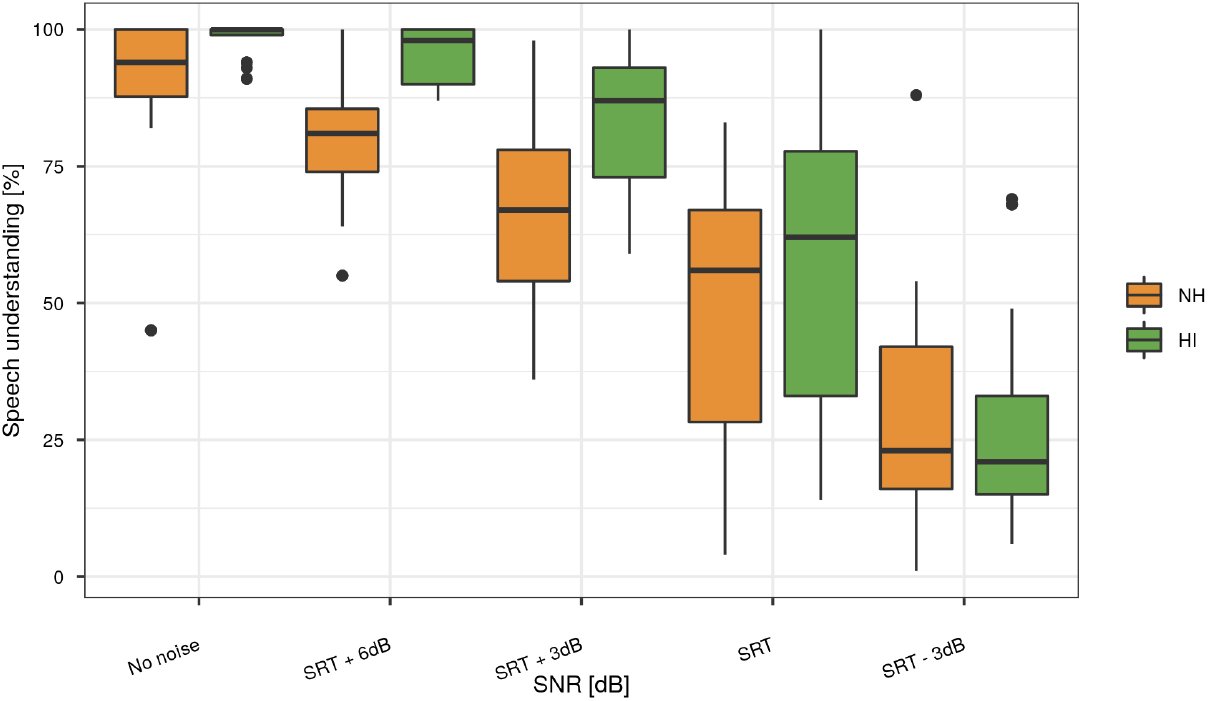
Subjective rating for each SNR condition for hearing-impaired listeners (green) and normal hearing listeners (orange).

## Spatial Filters

**Figure S.2:**
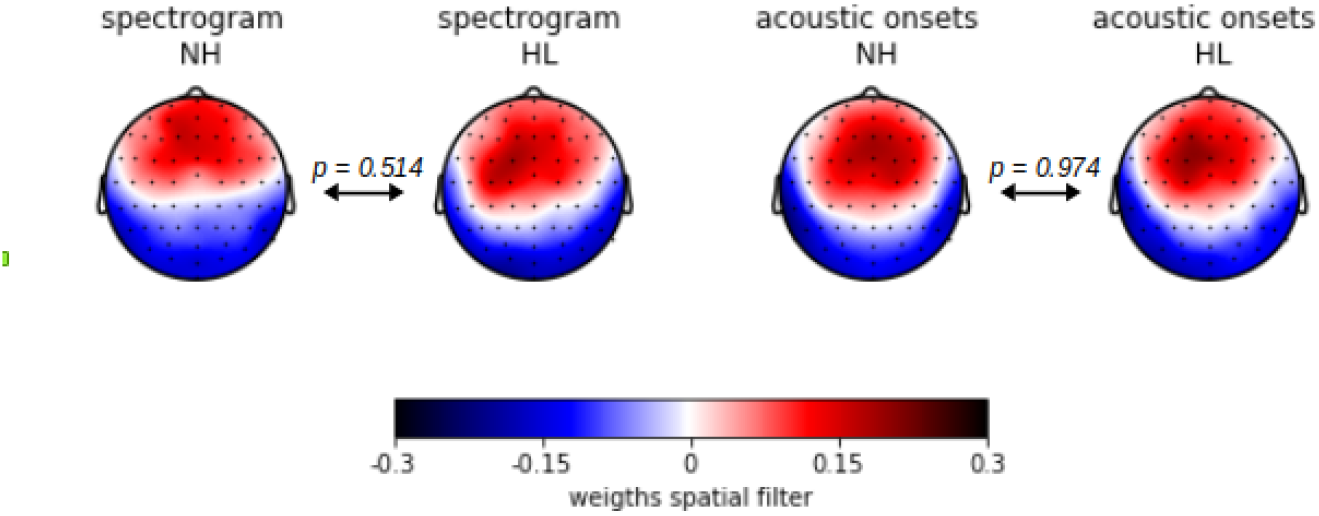
Spatial filters for acoustic onsets and spectrogram TRF, evaluated for both NH listeners and HI listeners.

## Supplementary Statistical Material

**Table S.1:** Results of the linear mixed model in order to assess the effects of degree of hearing loss (PTA) and age on the peak latency of *P*1_*AO*_, *N*1_*AO*_, *N*1_*S*_, *P*2_*S*_. Estimates of the regression coefficients (*β*), standard errors (SE), degrees of freedom (df), t-Ratios and p-values are reported per fixed effect term. Participant nested in the matching factor was included as a random nested effect. Formula: latency ~ 1 + peak + PTA +age + peak:age + peak:PTA + peak:PTA:age + (1 | match/participant)

| fixed effect term | $\beta$ | SE | df | t-Ratio | p-value |
| --- | --- | --- | --- | --- | --- |
| Intercept (for P1 - acoustic onsets) | 59.7545 | 5.7913 | 45.0463 | 10.318 | $p < 0.001$ |
| peak = N1 - acoustic onsets | 63.1639 | 7.1839 | 46.8945 | 8.7925 | $p < 0.001$ |
| peak = N1 - spectrogram | 32.1969 | 8.2213 | 48.341 | 3.9163 | $p < 0.001$ |
| peak = P2 - spectrogram | 141.4255 | 6.1896 | 42.5149 | 22.8488 | $p < 0.001$ |
| Degree of hearing loss (PTA) | 12.8954 | 3.5184 | 17.8724 | 3.6652 | $p = 0.0018$ |
| Age | -11.3602 | 9.4021 | 41.6263 | -1.2083 | $p = 0.2338$ |
| peak = N1 - acoustic onsets:age | 14.8838 | 11.4562 | 45.5116 | 1.2992 | $p = 0.2004$ |
| peak = N1 - spectrogram:age | 18.3418 | 12.275 | 46.5449 | 1.4942 | $p = 0.1419$ |
| peak = P2 - spectrogram:age | 32.7614 | 9.9271 | 41.8241 | 3.3002 | $p = 0.0020$ |

**Table S.2:** Results of the linear model in order to assess the effect of degree of hearing loss (z-scored) on the peak latency of *N*1_*AO*_ when only HI listeners are takening into account. Estimates of the regression coefficients (*β*), standard errors (SE), t-Ratios and p-values are reported per fixed effect term. Formula: *N*1_*AO*_-latency ~ 1 + PTA *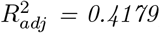, F = 8.178 on 1 and 9 df, p = 0.01879*

| fixed effect term | $\beta$ | SE | t-Ratio | p-value |
| --- | --- | --- | --- | --- |
| Intercept | 114.79 | 11.41 | 10.06 | $p < 0.001$ |
| Degree of hearing loss (PTA) | 34.89 | 12.20 | 2.86 | 0.0188 |

## References

Aiken, S. J. and Picton, T. W. (2008). Human cortical responses to the speech envelope. Ear and hearing, 29(2):139–157.

Alain, C. (2014). Effects of age-related hearing loss and background noise on neuromagnetic activity from auditory cortex. Frontiers in systems neuroscience, 8:8.

Bertoli, S., Probst, R., and Bodmer, D. (2011). Late auditory evoked potentials in elderly long-term hearing-aid users with unilateral or bilateral fittings. Hearing research, 280(1-2):58–69.

Bidelman, G. M., Mahmud, M. S., Yeasin, M., Shen, D., Arnott, S. R., and Alain, C. (2019a). Age-related hearing loss increases full-brain connectivity while reversing directed signaling within the dorsal–ventral pathway for speech. Brain Structure and Function, 224(8):2661–2676.

Bidelman, G. M., Price, C. N., Shen, D., Arnott, S. R., and Alain, C. (2019b). Afferent-efferent connectivity between auditory brainstem and cortex accounts for poorer speech-in-noise comprehension in older adults. Hearing research, 382:107795.

Billings, C. J., Penman, T. M., McMillan, G. P., and Ellis, E. (2015). Electrophysiology and perception of speech in noise in older listeners: effects of hearing impairment & age. Ear and hearing, 36(6):710.

Brodbeck, C. (2020). Eelbrain 0.32. http://doi.org/10.5281/zenodo.3923991.

Brodbeck, C., Hong, L. E., and Simon, J. Z. (2018). Rapid transformation from auditory to linguistic representations of continuous speech. Current Biology, 28(24):3976–3983.

Brodbeck, C., Jiao, A., Hong, L. E., and Simon, J. Z. (2020). Neural speech restoration at the cocktail party: Auditory cortex recovers masked speech of both attended and ignored speakers. bioRxiv, page 866749.

Brodbeck, C. and Simon, J. Z. (2020). Continuous speech processing. Current Opinion in Physiology.

Burkard, R. F., Eggermont, J. J., and Don, M. (2007). Auditory evoked potentials: basic principles and clinical application. Lippincott Williams & Wilkins.

Byrne, D., Dillon, H., Ching, T., Katsch, R., and Keidser, G. (2001). Nal-nl1 procedure for fitting nonlinear hearing aids: characteristics and comparisons with other procedures. Journal of the American academy of audiology, 12(1).

Campbell, J. and Sharma, A. (2013). Compensatory changes in cortical resource allocation in adults with hearing loss. Frontiers in systems neuroscience, 7:71.

Cardin, V. (2016). Effects of aging and adult-onset hearing loss on cortical auditory regions. Frontiers in Neuroscience, 10:199.

Coren, S. (1993). The lateral preference inventory for measurement of handedness, footedness, eyedness, and earedness: Norms for young adults. Bulletin of the Psychonomic Society, 31(1):1–3.

Das, N., Biesmans, W., Bertrand, A., and Francart, T. (2016). The effect of head-related filtering and ear-specific decoding bias on auditory attention detection. Journal of neural engineering, 13(5):056014.

David, S. V., Mesgarani, N., and Shamma, S. A. (2007). Estimating sparse spectro-temporal receptive fields with natural stimuli. Network: Computation in neural systems, 18(3):191–212.

Decruy, L., Das, N., Verschueren, E., and Francart, T. (2018). The self-assessed Békesy procedure: validation of a method to measure intelligibility of connected discourse. Trends in hearing, 22:2331216518802702.

Decruy, L., Vanthornhout, J., and Francart, T. (2019). Evidence for enhanced neural tracking of the speech envelope underlying age-related speech-in-noise difficulties. Journal of neurophysiology, 122(2):601–615.

Decruy, L., Vanthornhout, J., and Francart, T. (2020). Hearing impairment is associated with enhanced neural tracking of the speech envelope. Hearing Research, page 107961.

Di Liberto, G. M., O’Sullivan, J. A., and Lalor, E. C. (2015). Low-frequency cortical entrainment to speech reflects phoneme-level processing. Current Biology, 25(19):2457–2465.

Dimitrijevic, A., Smith, M. L., Kadis, D. S., and Moore, D. R. (2019). Neural indices of listening effort in noisy environments. Scientific Reports, 9(1):1–10.

Ding, N. and Simon, J. Z. (2012a). Emergence of neural encoding of auditory objects while listening to competing speakers. Proceedings of the National Academy of Sciences, 109(29):11854–11859.

Ding, N. and Simon, J. Z. (2012b). Neural coding of continuous speech in auditory cortex during monaural and dichotic listening. Journal of neurophysiology, 107(1):78–89.

Eggermont, J. J. (2017). Acquired hearing loss and brain plasticity. Hearing Research, 343:176–190.

Etard, O. and Reichenbach, T. (2019). Neural speech tracking in the theta and in the delta frequency band differentially encode clarity and comprehension of speech in noise. Journal of Neuroscience, 39(29):5750–5759.

Francart, T., van Wieringen, A., and Wouters, J. (2008). Apex 3: a multi-purpose test platform for auditory psychophysical experiments. Journal of neuroscience methods, 172(2):283–293.

Fuglsang, S. A., Märcher-Rørsted, J., Dau, T., and Hjortkjær, J. (2020). Effects of sensorineural hearing loss on cortical synchronization to competing speech during selective attention. Journal of Neuroscience, 40(12):2562–2572.

Hamilton, L. S., Edwards, E., and Chang, E. F. (2018). A spatial map of onset and sustained responses to speech in the human superior temporal gyrus. Current Biology, 28(12):1860–1871.

Harkrider, A. W., Plyler, P. N., and Hedrick, M. S. (2006). Effects of hearing loss and spectral shaping on identification and neural response patterns of stop-consonant stimuli. The Journal of the Acoustical Society of America, 120(2):915–925.

Harkrider, A. W., Plyler, P. N., and Hedrick, M. S. (2009). Effects of hearing loss and spectral shaping on identification and neural response patterns of stop-consonant stimuli in young adults. Ear and hearing, 30(1):31–42.

Heeris, J. (2014). Gammatone filterbank toolkit 1.0. https://github.com/detly/gammatone.

Horton, C., Srinivasan, R., and D’Zmura, M. (2014). Envelope responses in single-trial eeg indicate attended speaker in a ‘cocktail party’. Journal of neural engineering, 11(4):046015.

Iotzov, I. and Parra, L. C. (2019). EEG can predict speech intelligibility. Journal of Neural Engineering, 16(3):036008.

Koerner, T. K. and Zhang, Y. (2018). Differential effects of hearing impairment and age on electrophysiological and behavioral measures of speech in noise. Hearing research, 370:130–142.

Kraus, F., Tune, S., Ruhe, A., Obleser, J., and Woestmann, M. (2020). Unilateral acoustic degradation delays attentional separation of competing speech. bioRxiv.

Lenth, R. (2020). emmeans: Estimated Marginal Means, aka Least-Squares Means. R package version 1.4.8.

Lesenfants, D., Vanthornhout, J., Verschueren, E., Decruy, L., and Francart, T. (2019). Predicting individual speech intelligibility from the cortical tracking of acoustic-and phonetic-level speech representations. Hearing research, 380:1–9.

Luts, H., Jansen, S., Dreschler, W., and Wouters, J. (2014). Development and normative data for the Flemish/Dutch matrix test.

Maamor, N. and Billings, C. J. (2017). Cortical signal-in-noise coding varies by noise type, signal-to-noise ratio, age, and hearing status. Neuroscience letters, 636:258–264.

Margolis, R. H. and Saly, G. L. (2008). Asymmetric hearing loss: definition, validation, and prevalence. Otology & Neurotology, 29(4):422–431.

Maris, E. and Oostenveld, R. (2007). Nonparametric statistical testing of eeg-and meg-data. Journal of neuroscience methods, 164(1):177–190.

Mateos-Aparicio, P. and Rodríguez-Moreno, A. (2019). The impact of studying brain plasticity. Frontiers in cellular neuroscience, 13:66.

McClannahan, K. S., Backer, K. C., and Tremblay, K. L. (2019). Auditory evoked responses in older adults with normal hearing, untreated, and treated age-related hearing loss. Ear and hearing, 40(5):1106–1116.

Mirkovic, B., Debener, S., Schmidt, J., Jaeger, M., and Neher, T. (2019). Effects of directional sound processing and listener’s motivation on eeg responses to continuous noisy speech: Do normal-hearing and aided hearing-impaired listeners differ? Hearing Research, 377:260–270.

Nasreddine, Z. (2004). Montreal cognitive assessment (MoCA). École des sciences de la réadaptation, Sciences de la santé, Université d’Ottawa.

Oates, P. A., Kurtzberg, D., and Stapells, D. R. (2002). Effects of sensorineural hearing loss on cortical event-related potential and behavioral measures of speech-sound processing. Ear and hearing, 23(5):399–415.

Oostenveld, R. and Praamstra, P. (2001). The five percent electrode system for high-resolution eeg and erp measurements. Clinical neurophysiology, 112(4):713–719.

O’Sullivan, J. A., Power, A. J., Mesgarani, N., Rajaram, S., Foxe, J. J., Shinn-Cunningham, B. G., Slaney, M., Shamma, S. A., and Lalor, E. C. (2015). Attentional selection in a cocktail party environment can be decoded from single-trial eeg. Cerebral cortex, 25(7):1697–1706.

Peelle, J. E. and Wingfield, A. (2016). The neural consequences of age-related hearing loss. Trends in neurosciences, 39(7):486–497.

Petersen, E. B., Wöstmann, M., Obleser, J., and Lunner, T. (2017). Neural tracking of attended versus ignored speech is differentially affected by hearing loss. Journal of neurophysiology, 117(1):18–27.

Presacco, A., Simon, J. Z., and Anderson, S. (2016). Evidence of degraded representation of speech in noise, in the aging midbrain and cortex. Journal of neurophysiology, 116(5):2346–2355.

Presacco, A., Simon, J. Z., and Anderson, S. (2019). Speech-in-noise representation in the aging midbrain and cortex: Effects of hearing loss. PloS one, 14(3):e0213899.

R Core Team (2020). R: A Language and Environment for Statistical Computing. R Foundation for Statistical Computing, Vienna, Austria.

Slaney, M. (1998). Auditory toolbox. Interval Research Corporation, Tech. Rep, 10(1998).

Somers, B., Francart, T., and Bertrand, A. (2018). A generic eeg artifact removal algorithm based on the multi-channel wiener filter. Journal of neural engineering, 15(3):036007.

Tremblay, K. L., Piskosz, M., and Souza, P. (2003). Effects of age and age-related hearing loss on the neural representation of speech cues. Clinical Neurophysiology, 114(7):1332–1343.

Van Dun, B., Kania, A., and Dillon, H. (2016). Cortical auditory evoked potentials in (un) aided normal-hearing and hearing-impaired adults. In Seminars in hearing, volume 37, page 9. Thieme Medical Publishers.

Vanthornhout, J., Decruy, L., Wouters, J., Simon, J. Z., and Francart, T. (2018). Speech intelligibility predicted from neural entrainment of the speech envelope. Journal of the Association for Research in Otolaryngology, 19(2):181–191.

Verschueren, E., Vanthornhout, J., and Francart, T. (2021). The effect of stimulus intensity on neural envelope tracking. Hearing Research, 403:108175.

Voeten, C. C. (2020). buildmer: Stepwise Elimination and Term Reordering for Mixed-Effects Regression. R package version 1.6.

